# Host avoidance and resistance vary independently and are specific to parasite genotype

**DOI:** 10.1101/2025.01.08.631927

**Authors:** Caroline R. Amoroso, Adianna Lockwood-Shabat, Andreas Kamali, Amanda K. Gibson

## Abstract

Hosts can reduce the negative fitness effects of parasite infection by avoiding contact with parasites or by resisting infection after contact. Because of their shared outcome, avoidance and resistance have been hypothesized to trade off with one another. Assuming these defenses carry fitness costs, hosts are expected to have high levels of one defense or the other, but not both. Alternatively, avoidance and resistance may covary positively, if, for example, they complement one another or are genetically or mechanistically linked. Testing these hypotheses requires measuring avoidance and resistance independently, which is challenging because they are functionally linked. In this study, we separated avoidance and resistance of the host *C. elegans* against the bacterial parasite *S. marcescens* and tested for a correlation between them. We phenotyped a panel of 12 genetically divergent hosts using two distinct bacterial strains and multiple experimental contexts. We found no evidence of a correlation between avoidance and resistance. This result suggests that avoidance and resistance can covary independently. Moreover, we found strong genetic specificity not only for resistance, but also for a measure of avoidance, motivating further research to examine the coevolutionary dynamics of avoidance.

## Introduction

Defense against parasites can take many forms. Hosts can avoid parasite infection by reducing their contact rate with parasites, or they can resist infection by reducing parasite establishment and/or proliferation after contact has been made [1]. Avoidance and resistance have similar functions: they both increase host fitness in the presence of parasites and reduce parasite fitness. Their shared effects make it challenging to quantify their distinct contributions to infection outcomes. Controlled experiments are required to independently measure avoidance and resistance. Furthermore, understanding each defense’s independent role is complicated by the potential for them to covary as a result of natural selection and/or other evolutionary and population genetic processes [2]. Thus, to understand variation in defense across individual hosts, it is important to examine the covariance between independent measures of avoidance and resistance in a genetically explicit way that identifies the possibility for these traits to evolve jointly. The goal of this study is to determine the direction and genetic basis of the covariance between avoidance and resistance.

The dominant hypothesis in the literature is that selection should cause avoidance and resistance to covary negatively. If avoidance and resistance are both costly, then individuals should be selected to allocate energy or resources to one or the other defense, but not both [3– 7]. This hypothesis assumes that avoidance and resistance are functionally redundant, such that having both does not provide greater protection than having one. In several studies of birds, avoidance of sick-behaving conspecifics covaries negatively with proxies for immune function across individuals [3–5]. In trout and salmon, avoidance and resistance did not covary across individuals [6], but covaried negatively across populations [7]. These studies support a negative covariance of avoidance and resistance, raising the question of the genetic basis of these traits.

Alternatively, selection could generate a positive covariance between avoidance and resistance. We expect this if avoidance and resistance are complementary, rather than redundant, such that having both avoidance and resistance provides greater protection than having just one. In this case, selection could favor higher levels of both traits. Supporting this prediction, families of white campion (*Silene latifolia*) that phenologically avoided a fungal parasite also had higher resistance when directly inoculated [8]. *Caenorhabditis elegans* populations evolved both elevated avoidance and resistance over 30 generations of exposure to the parasite *Serratia marcescens* [9,10]. Linkage could also generate a positive covariance. For example, the mechanisms of avoidance and resistance of *C. elegans* to a parasitic strain of *Bacillus thuringiensis* both seem to involve direct detection of the parasite and so covary positively [11]. There is also some evidence in humans that triggering avoidance behaviors can increase production of immune cells [12,13]. Linkage could also be genetic, although this has not been demonstrated empirically to our knowledge.

In this study, we measure the covariance between avoidance and resistance in the nematode host *C. elegans* and its bacterial parasite *S. marcescens. C. elegans* strains are known to vary in avoidance and resistance to *S. marcescens* [14,15], but it is not known whether there is any relationship between the two defenses. We focus on 12 genetically divergent strains of *C. elegans* to capture a broad range of genetic variation in the two defenses. We independently measure the host strains’ avoidance and resistance to two distinct strains of *S. marcescens*: one known to be highly virulent [15,16], and the other highly avoided [14,17]. We examine whether there are correlations between avoidance and resistance traits across *C. elegans* strains, as well as between composite measures of avoidance and resistance.

## Methods

### Overview

We examined the relationship between avoidance and resistance to two distinct strains of the opportunistic bacterial parasite *S. marcescens* across 12 genetically divergent strains of *C. elegans*. We chose parasite strains that we expected to differ in their virulence and the avoidance they elicit. We assayed resistance independently of avoidance by measuring survival after directly dosing hosts in liquid, which prevents hosts avoiding the parasite. We used two assays to measure avoidance: a lawn-leaving assay [17] and a choice assay [11]. These are two common designs used for assessing behavioral responses to bacteria, and we expected they would capture different aspects of behavioral variation. We used data from these assays to test whether resistance and avoidance covaried across *C. elegans* strains. Each assay was conducted in a blocked design, with a subset of host strains tested in each of four to five experimental blocks that contained all treatments (see Table S1 for block assignments and Supplementary Text – Section S1 for details on statistical handling of block effects).

### Strains

#### Hosts

The 12 genetically divergent strains of *C. elegans* were acquired from the *Caenorhabditis* Natural Diversity Resource (CaeNDR, https://caendr.org/). The strains were as follows: CB4856, CX11314, DL238, ED3017, EG4725, JT11398, JU258, JU775, LKC34, MY16, MY23, and N2. (We assigned host strains a number label from 1 to 12 to improve interpretation across the figures; see Table S1.) CaeNDR recommends using this set of strains as a starting point to determine whether genetic variation exists in ecologically relevant traits across the species [18]. In a previous study, these strains showed heritable variation in avoidance behavior (lawn-leaving, see below) of the *S. marcescens* strain Db10 [14].

Upon receipt of the strains, we followed the CaeNDR protocols for cleaning the strains (see https://caendr.org/data/protocols), and we cryopreserved them at −80°C. Before experiments, we thawed the strains and maintained them on ample food (*Escherichia coli* strain OP50) for 14–17 days at 20°C prior to an assay to allow populations to recover from freezing.

#### Parasites

The two *S. marcescens* strains used in this experiment were Sm2170 and Db10. Sm2170 is a red-pigmented strain that is known to be highly virulent, killing the greatest proportion of hosts among a set of five *S. marcescens* strains in a previous study [15]. Prior studies found that this parasite strain is sufficiently virulent to select for avoidance and resistance in the host [9,10,16,19]. The second strain, Db10, is unpigmented and elicits a high degree of avoidance from *C. elegans* [14,17]. The virulence of Db10 and Sm2170 has not been directly compared, but Db10 appears to be less virulent [17]. Stocks of the bacterial strains were maintained at - 80ºC in glycerol and thawed for proliferation prior to each experiment.

### Experimental Design

#### Quantifying Resistance

To isolate resistance (post-contact defenses) from avoidance (pre-contact), we measured infection outcomes after controlling for contact, i.e., by directly dosing the host with a known concentration of bacterial cells. We accomplished this in a liquid environment, where hosts indiscriminately ingested a suspension of the parasite. Then we monitored survival over the subsequent days [11,15]. We used two doses of each of the two parasite strains to maximize the potential variation in resistance that we could capture across the 12 host strains (Fig. 1A).

**Figure 1.**
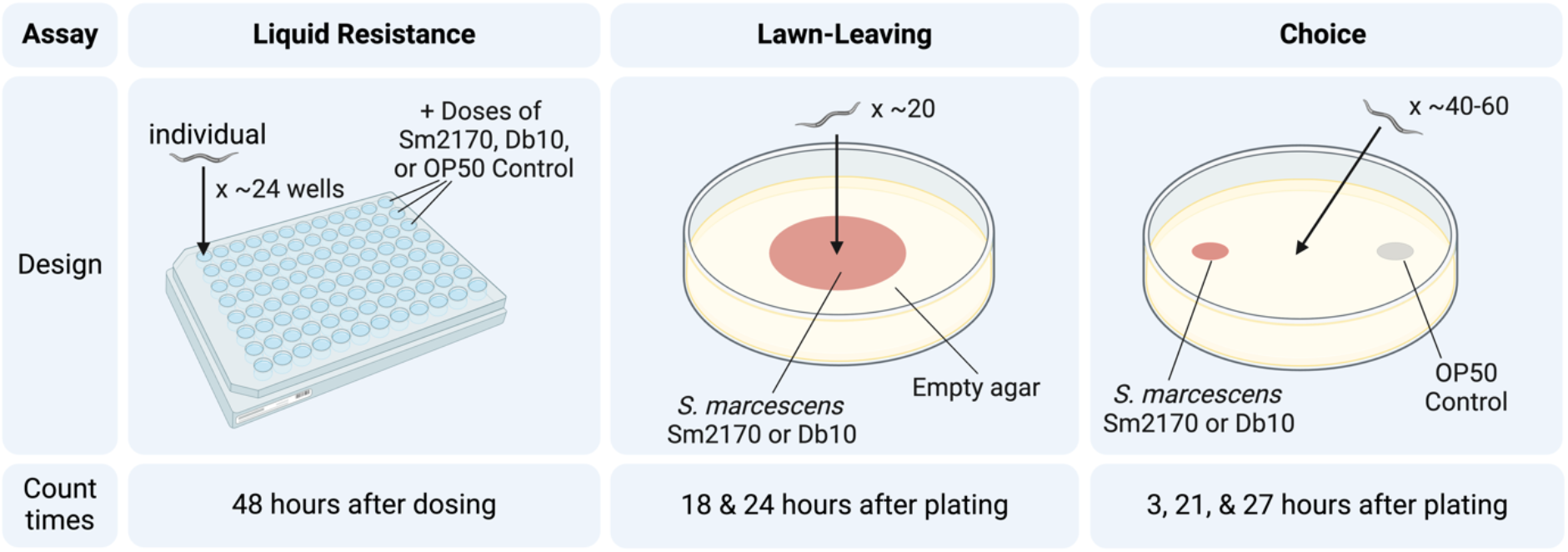
Experimental designs for the three defense assays. Figure created with BioRender.com.

For each *C. elegans* strain, L4 larval stage nematodes were individually sorted into wells of a 96-well plate using a Union Biometrica Large Particle Flow Cytometer (WormSorter) that initially contained 50 µl of sterile S-medium. Each well was inspected microscopically to confirm the presence of one live nematode. We excluded wells with more than one nematode or a dead nematode. Then the hosts were dosed with an additional 50 µl of one of five treatments: a high or a low dose of Sm2170, a high or a low dose of Db10, or a control dose of OP50. The high and low doses were prepared by pelleting 24-hour LB cultures of the two parasite strains (initial bacterial concentrations: Sm2170 4.45 × 10^8^ CFU/ml; Db10 4.73 × 10^8^ CFU/ml) in a centrifuge and resuspending them in one-eighth (high dose) or one-quarter (low dose) volume of S-medium. When added to the 50 µl of S-medium already in the wells, the high dose was concentrated 4x from the initial culture’s cell counts, and the low dose was concentrated 2x. For the OP50 control, a 24-hour culture in LB was pelleted and resuspended in an equal volume of S-medium. Because it included no nutrients, S-medium did not promote additional growth of the bacteria in the wells over the course of the experiment. Host strains were assigned to wells column-wise in a balanced design across four to five plates, and bacterial strain doses were administered row-wise (see Figure S1 for details). For each combination of host strain, parasite strain, and treatment, the WormSorter filled on average 28.8 +/-7.45 wells with a single L4-stage host. Each plate was covered with Breathe-Easy film and incubated at 20ºC, shaking at 160 rpm to promote gas exchange in the liquid.

After 48 hours, we inspected each well with an inverted microscope and scored individual hosts as alive if there was any sign of movement [11]. We also recorded whether larvae were present in the well, indicating that the host had successfully reproduced. Individuals that were missing at 48 hours were excluded. We obtained survival data for a mean of 27.6 +/-7.5 individuals per combination of host strain, parasite strain, and treatment, ranging from a minimum of 10 to a maximum of 46 individuals per combination.

#### Quantifying Avoidance with a Lawn-leaving Assay

We first measured avoidance using a lawn-leaving assay (Fig. 1B), which is modeled after past studies of avoidance of strain Db10 [14,17]. We prepared the lawns by pipetting 30µL of 6-hour LB cultures of Db10 or Sm2170 onto the center of 60-mm Petri dishes of nematode growth medium (NGM-Lite, US Biological, Catalog No: N1005) with 0.7% agarose added to discourage *C. elegans’* burrowing. Using an inoculating loop, lawns were spread to cover a 25-mm-diameter circle that had been drawn on the bottom of the Petri dish with a template to standardize the lawn’s size, shape, and location. The plates were incubated overnight at 28ºC.

Synchronized populations of nematodes were raised on OP50 for 48 hours at 20ºC until they reached the L4 larval stage. They were washed from the plates and into a tube with M9. They were washed three times with 2 ml of M9 to remove OP50, and then approximately 20 individuals were transferred in ∼3 µl of M9 to the center of the bacterial lawn using a pipette. The precise number of hosts added was immediately confirmed by microscopy.

After hosts were added, the plates were maintained at 20ºC. After 18 and 24 hours, we counted the number of individuals located on and off the lawn of bacteria. Nematodes touching any part of the lawn were counted as on the lawn. The baseline expectation for non-parasitic bacteria is that nearly all individuals would be located on the lawn of bacteria at this timepoint, as demonstrated by the control treatment in a previous study [14]. This experimental protocol follows that of the previously published study [14], except that the transfer of nematodes was done in liquid, which is more efficient than picking individual nematodes.

#### Quantifying Avoidance with a Choice Assay

We also measured behavioral responses to *S. marcescens* in a binary choice assay (Fig. 1C), which is a common experimental design for studying *C. elegans*’ bacterial preference. Here, we grew cultures of Db10 and Sm2170 in LB for 6 hours. We also grew a culture of OP50 overnight. Onto each 100-mm Petri dish containing nematode growth medium with 0.7% agarose, we added two 10-µl spots of bacterial culture: one spot of OP50, and the other spot of either Db10 or Sm2170. The bacterial spots were added on opposite sides of the dish, each approximately 1 cm from the inner wall of the dish. The dishes were incubated overnight at 28ºC to grow into small lawns.

*C. elegans* populations were prepared as for the lawn-leaving assay, and approximately 50 L4-stage nematodes were added to the center of the agar, equidistant from the two bacterial spots. An initial count of nematodes was recorded within the first 10 minutes of plating. The number of nematodes located on each lawn and on neither lawn was counted at 3 hours, 21 hours, and 27 hours after plating.

### Data Analysis

Analyses were run in R (v. 4.4.1) [20] using RStudio (v 2024.04.2+764) [21]. Data were processed and graphics produced with the package *tidyverse* [22].

We first used the liquid resistance data to examine the effects of each parasite strain on survival compared to the OP50 control. Survival was coded as binary (1/0). We fit a generalized linear mixed model (‘glmer’ function in the package *lme4* [23]) with a binomial family that included bacterial strain and *C. elegans* strain as fixed effects, and plate as a random effect. To examine variation in resistance to the parasites across host strains, we fit a second binomial GLMM to the data only for the parasite treatments, with host strain, parasite strain, and dose as fixed effects. We controlled for plate with a random effect.

We then used the avoidance data to examine variation across host and parasite strains. For the lawn-leaving data, we fit a binomial generalized linear mixed model to the counts of hosts on and off the parasite lawns, using host strain, parasite strain, and count time as predictors, and including a random effect for plate. For the choice assay, we calculated a ‘choice index’ for each plate by subtracting the number of hosts on the *S. marcescens* lawn from the number on the OP50 lawn and dividing by the total, including hosts on neither lawn. A negative choice index indicates attraction to the parasite, and positive choice index indicates avoidance. We fit a linear mixed model that included host strain, parasite strain, count time, and block as fixed effects, and plate as a random effect.

Model fits were assessed with the ‘check_model’ function and its extensions in the package *performance* [24] and tested for overdispersion where appropriate. We estimated marginal means (‘emmeans’ function from the package *emmeans* [25]) and where appropriate, converted them from log-odds to percentages for ease of interpretation (‘plogis’ function). Models were compared using AIC scores (‘AIC’ and ‘compare_performance’ functions).

We took two approaches to examine correlations between avoidance and resistance. The first was to run pairwise correlations (‘cor’ function) between all the avoidance and resistance trait variables to determine whether any individual measures of avoidance and resistance were correlated. We performed a Bonferroni adjustment to correct for multiple comparisons. The second was to conduct a principal components analysis (PCA) on the avoidance traits and a separate PCA on the resistance traits to capture a single axis of variation within each defense category (‘princomp’ function). We then fit a linear model to the major axis for avoidance and the major axis for resistance. For strain-level estimates of each defense trait, we took the mean trait value across replicates for each host strain for each treatment, and logit-transformed the variables that were measured as proportions.

## Results

### Variation in Resistance

Across all *C. elegans* strains, the estimated marginal mean survival was 45.6% (CI_95_=[40.1, 51.3]) for Sm2170 and 84.0% (CI_95_=[79.9, 87.3]) for Db10, compared to 98.0% (CI_95_=[96.6, 98.9]) for the OP50 control. Thus both *S. marcescens* strains were detrimental to *C. elegans* survival, and Sm2170 caused greater mortality than Db10.

Comparing survival across the host and two parasite strains, the model that was best supported by the data included a three-way interaction between parasite strain, host strain, and dose (ΔAIC>31; Table S2). Thus, host strains varied in resistance, and host strains that were more resistant to one strain of the parasite were not necessarily more resistant to the other (Fig. 2). The effect of dose also depended on the specific host-parasite pairing. These results reflect a high degree of specificity in the host-parasite interaction.

**Figure 2:**
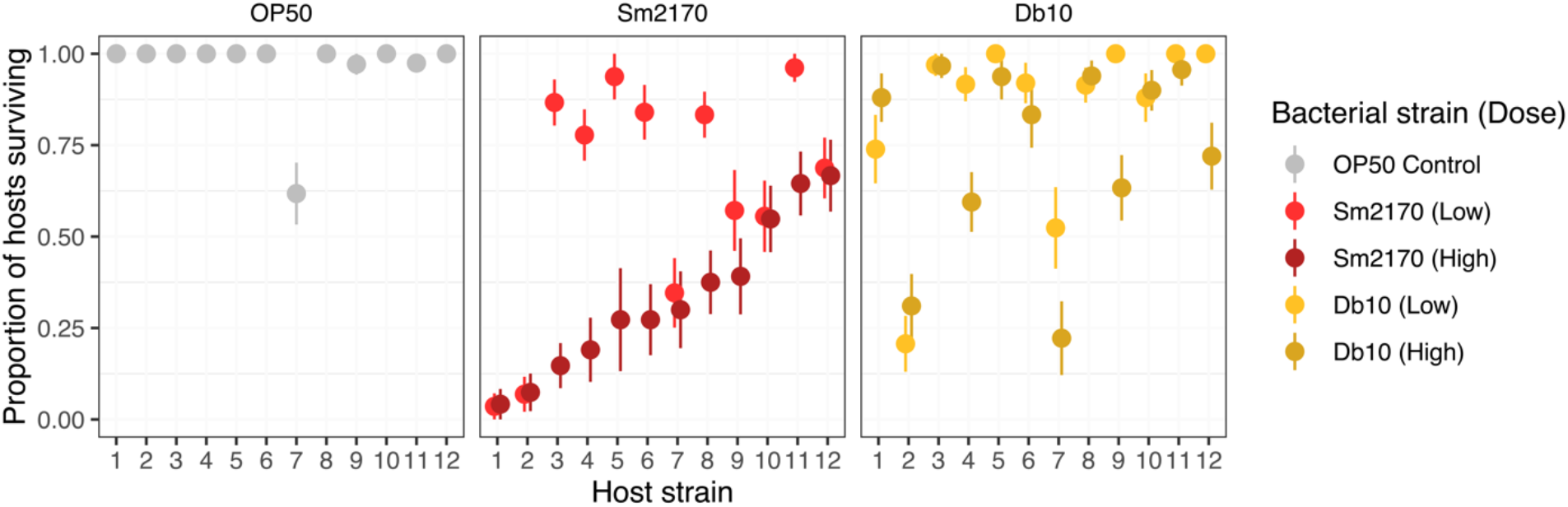
Resistance Variation. Survival of 12 *C. elegans* host strains in two doses of two strains of the bacterial parasite *S. marcescens* and an OP50 control. Survival was recorded after 48 hours of continuous exposure in liquid, preventing avoidance behavior and thereby isolating post-contact defense. *C. elegans* strains are ordered on the axis by survival in the high dose of Sm2170. Points and error bars represent means and standard errors.

**Figure 3:**
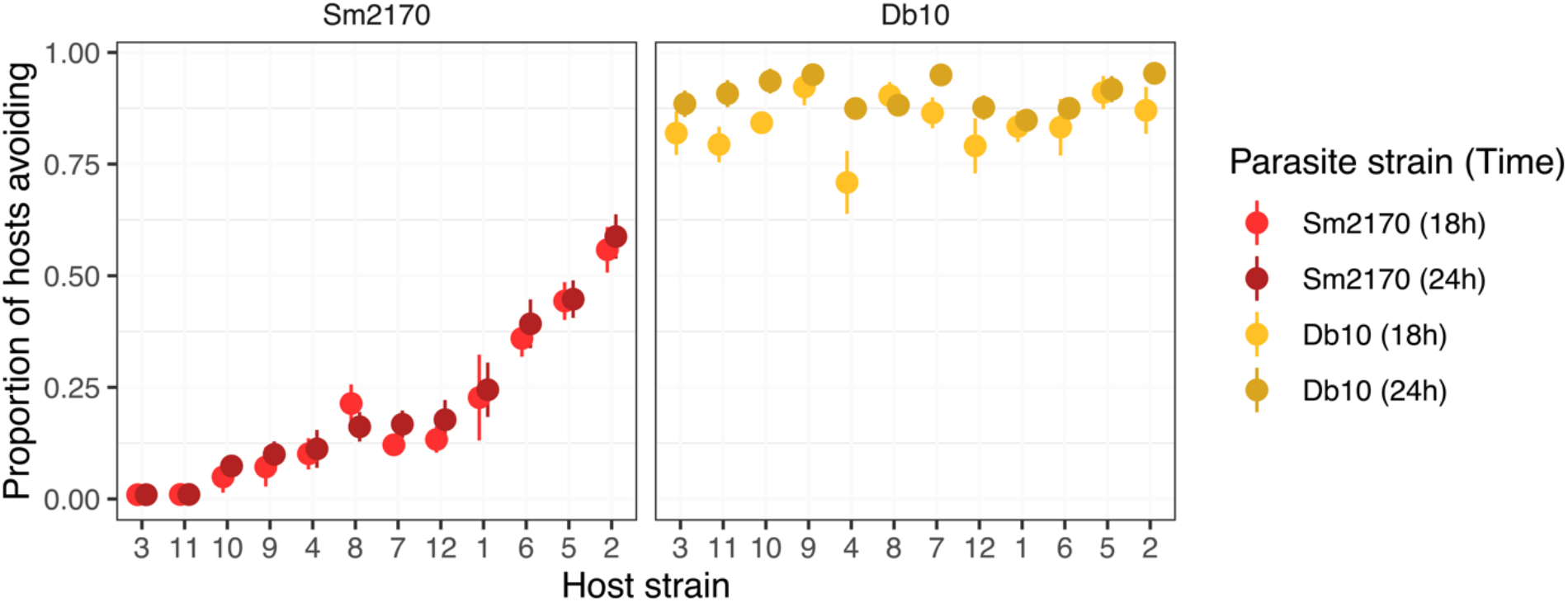
Lawn-leaving variation. Host strains vary in the proportion of individuals that leave lawns of two strains of the parasite *S. marcescens. C. elegans* host strains are ordered on the axis by their avoidance of Sm2170 at 24 hours. Points and error bars represent means and standard errors.

Both *S. marcescens* strains also reduced host fecundity relative to the *E. coli* control. The marginal mean percentage of individuals producing larvae at 48 hours in the OP50 control was 95.1% (CI_95_=[91.0, 97.4]). For Sm2170, it was 4.2% (CI_95_=[2.4, 7.1]), and for Db10, it was 17.2% (CI_95_=[11.1, 25.7]). The best supported model of fecundity included the three-way interaction between host strain, parasite strain, and dose (ΔAIC>6; Table S3). Thus, as for survival, effects of the parasite on host fecundity varied with dose and the specific interaction of host and parasite strain.

We noticed a paralytic effect of the high dose of Db10 when hosts were checked at 24 hours. Some hosts appeared immobile when assessed at 24 hours but were mobile again at 48 hours (Figure S2).

### Variation in Lawn-Leaving Avoidance Behavior

The two parasite strains produced distinct lawn-leaving responses. Across both timepoints and the dozen host strains, only 14.6% (CI_95_=[12.0, 17.7]) of individuals avoided the lawn of Sm2170, whereas 89% avoided the Db10 lawn (CI_95_=[86.7, 91.1]). Thus, there was much greater avoidance of Db10 than Sm2170 overall, and both strains were avoided more than an OP50 control reported for these same dozen strains in a previous study (2.9%; CI_95_=[2.1, 4.0]) [14]. The model that was best supported by the data included an interaction between host strain and parasite strain (ΔAIC>22; Table S4), indicating that host-parasite pairings were consistent across the time points.

### Variation in Choice Behavior

In the choice assay, *C. elegans* strains were initially attracted to the parasite, as indicated by a negative choice index at the 3-hour mark. By the 21- and 27-hour counts, strains overall showed more avoidance, with a greater relative proportion of individuals on the OP50 control lawn than the lawn of either *S. marcescens* strain. For strain Sm2170, the estimated marginal mean choice index at 3 hours was -0.12, and it increased to 0.09 and 0.10 at 21 and 27 hours, respectively. For Db10, marginal mean choice index was -0.07, 0.14, and 0.15 at the same time points, respectively. The 95% confidence intervals of these overall mean estimates for each parasite strain overlap at each timepoint, reflecting similar mean levels of avoidance across the two parasite strains. In the model comparison, we found that the best model included host strain, parasite strain, and count time with no interactions among them (ΔAIC>10; Table S5).

### Correlations among the defenses

The high degree of specificity we observed in some of the experiments indicated that we should consider correlations between the defense traits separately for the two parasite strains. We detected few significant pairwise correlations *between* avoidance and resistance measures (strain-level means) (Fig. 5). We detected marginally significant negative correlations between low dose survival and the choice index for Sm2170, as well as between high dose survival and lawn-leaving at 24 hours for Db10 (Fig. 5). However, after correcting for multiple comparisons, neither of these correlations reached statistical significance. Within avoidance and within resistance, trait correlations were generally positive, but most did not reach statistical significance when adjusting for multiple comparisons. There were two significant pairwise correlations within avoidance measures: between lawn-leaving at 18 hours and at 24 hours for Sm2170, and choice at 21 and 27 hours for Db10.

We performed PCA to estimate each host strain’s overall level of resistance and avoidance to each parasite strain that accounted for the covariance of traits within these defense categories. For resistance traits, we conducted a PCA on survival and fecundity at the high and low doses. For Sm2170, the first principal component (PC1) of the analysis accounted for 56.3% of the variance among host strains in resistance traits, and for Db10, it accounted for 67.8%. For avoidance traits, we conducted a PCA on data from the lawn-leaving assay at 18 and 24 hours and the choice assay at 21 and 27 hours. We opted to exclude the 3-hour choice assay data because it indicated attraction rather than avoidance (Fig. 4). In the Sm2170 analysis, PC1 accounted 57.5% of the variance among host strains in avoidance traits, and for Db10, PC1 accounted for 54.0%. We did not find a correlation between PC1 for avoidance and PC1 for resistance for either parasite strain (Fig. 6; Table S6).

**Figure 4:**
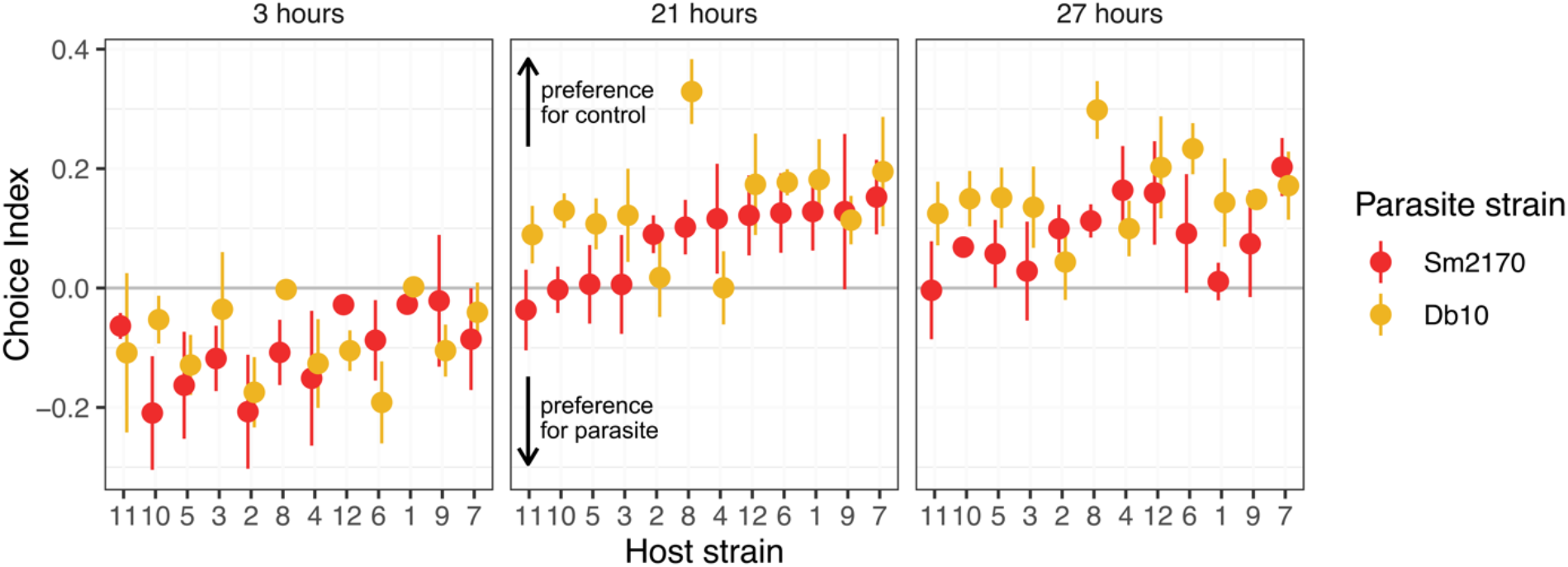
Choice variation. When given a choice between lawns of *S. marcescens* and *E. coli* strain OP50, *C. elegans* strains varied in their choice. The choice index is the proportion of total host individuals counted on the control OP50 lawn minus the proportion of total counted on the *S. marcescens* lawn. A positive choice index indicates a greater relative proportion of hosts on the OP50 lawn than on the *S. marcescens* lawn, and a negative choice index indicates a greater relative proportion on the *S. marcescens* lawn than the OP50. Panels show data for different amounts of time since plating hosts, and host strains are ordered on the axis by increasing avoidance of Sm2170 at the 21-hour time point. Points and error bars represent means and standard errors.

**Figure 5.**
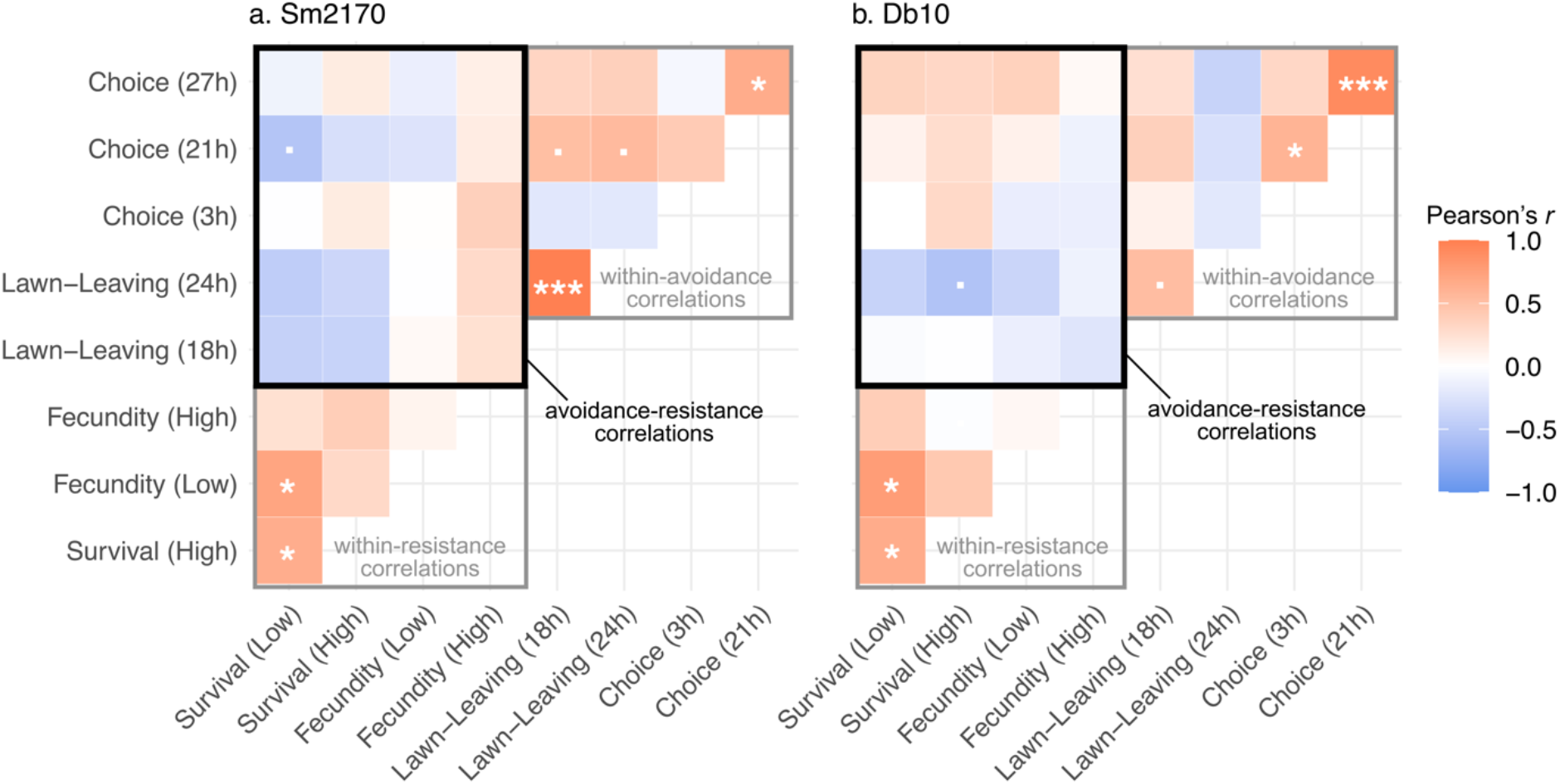
Defense trait correlations. Heatmaps showing pairwise correlations between each defense measurement across the 12 host strains, where the color and intensity of the shading corresponds to the Pearson’s correlation coefficient (*r*), and symbols represent significance levels before correcting for multiple comparisons (· p= <0.1; * p<0.05; *** p<0.001). The correlations denoted by *** are the only ones that are still statistically significant following Bonferroni adjustment. Correlations of each variable with itself (i.e., r=1 across the diagonal) are not included.

**Figure 6.**
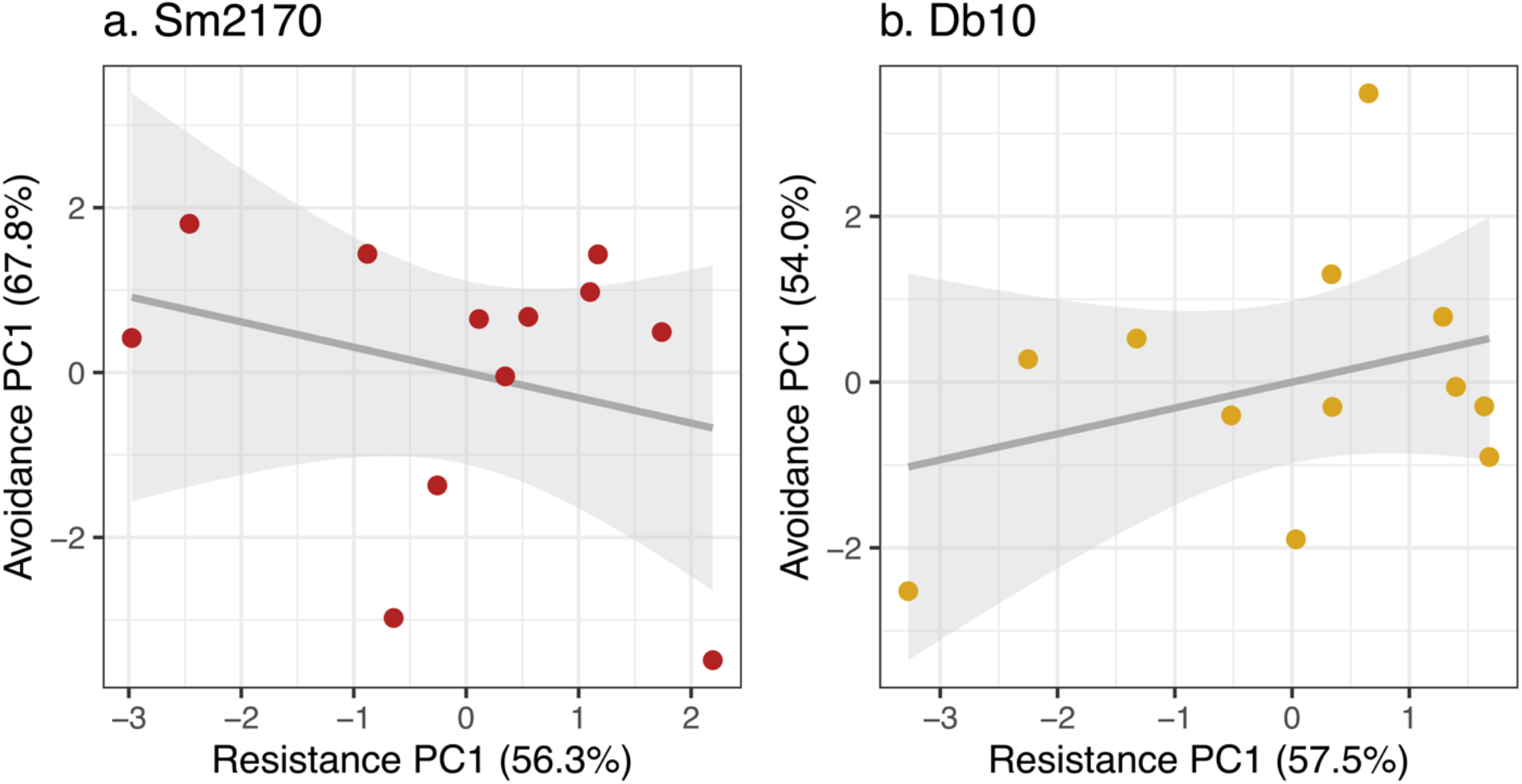
No correlation between avoidance and resistance first principal components. Each PC1 represents a main axis of variation across four covarying measures of avoidance and resistance to each parasite strain across 12 host strains. The number in parentheses represents the percent of the variance among host strains that is explained by the first principal component (PC1). There is no correlation between avoidance and resistance to Sm2170 (a) or Db10 (b).

## Discussion

This study captured an extensive range of variation in avoidance and resistance across a dozen genetically divergent host strains and two distinct parasite strains. Mean levels of both avoidance and resistance against Sm2170 were lower than against Db10. Within a parasite strain, there was no relationship between avoidance and resistance across host strains. This was true for pairwise correlations between individual avoidance and resistance measures, as well as for correlations across composite measures of each defense. In addition, we found evidence of highly specific strain-by-strain (GxG) effects for resistance and one of our avoidance measures (lawn-leaving). On the other hand, the second avoidance measure (choice) varied across host and parasite strains, but without evidence of a specific genetic interaction.

The absence of a relationship between avoidance and resistance in this study contrasts with previous research in several species of birds [3–5] and fish [6,7] that found evidence of a negative covariance between avoidance and resistance. The negative relationship between the defenses is hypothesized to result from their redundancy and the defenses having constitutive costs on host fitness. *C. elegans* has previously been shown to incur no constitutive costs of defense against *S. marcescens* [14,19], and this study is consistent with these prior findings. If avoidance and resistance are not costly, then the trade-off hypothesis may not apply.

The absence of a relationship between avoidance and resistance also contrasts with past research in *C. elegans*. A past study found a positive correlation between avoidance and resistance defenses to a different parasite species, *Bacillus thuringiensis*, which the authors hypothesized represented a mechanistic link between avoidance and resistance [11]. We did not find evidence of such a positive covariance among natural host strains in this study, rejecting such a link between avoidance and resistance. These results also contrast with an experimental evolution study in which *C. elegans* evolved both higher avoidance and resistance to *S. marcescens* over generations of selection [9,10]. A key difference of this study is that we do not know the history of these strains’ exposure to *S. marcescens* in the habitats from which they were sampled, although it is likely that *C. elegans* encounters *Serratia* spp. in its natural environment [26]. The specificity that we observed in this study suggests that selection by other strain(s) of *Serratia* spp. in the hosts’ natural environments would not necessarily translate to equivalent trait levels against the two *S. marcescens* strains we tested here.

We conducted two different avoidance assays to capture multiple potential axes of behavioral variation, because it was not clear what was the most relevant experimental context. There were neither strong nor consistent correlations between the lawn-leaving and the choice assays. The absence of a correlation is consistent with previous studies suggesting lawn-leaving and choice behaviors rely on distinct mechanisms [17,27]. The variation we detected within and across avoidance is consistent with the context-dependence of behavioral defenses.

Avoidance and resistance both showed a high degree of specificity in this study, with the best-supported models for both the liquid resistance assay and the lawn-leaving assay including a host-strain by parasite-strain interaction. Specificity is well-documented for resistance, and this specificity is key to the maintenance of genetic diversity in coevolving host and parasite populations [28]. In contrast, avoidance tends to be thought of as a more general defense [29]. Many well-described avoidance behaviors, such as feces avoidance [30–32], are likely to protect hosts from diverse parasite taxa that are transmitted via the same route. Our finding of highly specific genetic interactions underlying avoidance contrasts with this notion that avoidance is general. Thus, coevolutionary theory that relies upon specific host genotype-by-parasite genotype interactions may also apply to avoidance. Furthermore, it is striking that hosts had higher avoidance and resistance to Db10 than Sm2170 on average, suggesting a positive correlation between the defenses across parasite strains. These results motivate further testing against a broader array of parasite strains.

This study suggests that avoidance and resistance are not linked – mechanistically, genetically, or through constitutive fitness effects. However, their independent variance across diverse host strains does not exclude the possibility that avoidance and resistance could affect each other’s evolution. For example, natural selection acting upon avoidance and resistance in sequence could generate covariance between them [1,33]. By occurring first, avoidance may be under strong selection, and high levels of avoidance could block or prevent downstream selection by parasites on resistance. How these sequential dynamics would play out would depend on how the defense traits and their costs interact, as well as on resulting eco-evolutionary feedbacks [34].

## Supporting information

Amoroso et al. spplementary materials

## Acknowledgements

We thank Tessa Batterton for her helpful assistance with the resistance assays. We appreciate LTC Andrew Kick’s logistical support of the choice experiments. We would like to thank Dr. Janis Antonovics and members of the Gibson Lab for feedback on the project. The *E. coli* strain OP50 and *S. marcescens* strain Db10 used in this study were provided by the *Caenorhabditis* Genetics Center (CGC) at the University of Minnesota. The *S. marcescens* strain Sm2170 was provided by Dr. Levi Morran at Emory University.

