## Supplementary material for "Host avoidance and resistance vary independently and are specific to parasite genotype": Amoroso et al. spplementary materials

**Table S1: Strain assignments to experimental blocks**

Each of the 12 host strains was assigned a numeric label, making the figures in the main text more legible/interpretable. Each assay measured the defense level of a set of strains in 4-5 blocks, which were performed in different weeks but were otherwise identical. For the resistance and lawn-leaving assays, each strain was included in a single block, whereas for the choice assay, each strain was replicated across two blocks.

| **Host strain** | **Label** | **Resistance block** | **Lawn-leaving block** | **Choice blocks** |
| --- | --- | --- | --- | --- |
| CB4856 | 6 | 1 | 2 | 2, 4 |
| CX11314 | 7 | 4 | 3 | 1, 3 |
| DL238 | 1 | 5 | 2 | 1, 4 |
| ED3017 | 9 | 4 | 4 | 2, 3 |
| EG4725 | 5 | 3 | 3 | 2, 4 |
| JT11398 | 3 | 2 | 3 | 1, 4 |
| JU258 | 12 | 4 | 4 | 1, 3 |
| JU775 | 10 | 5 | 5 | 2, 3 |
| LKC34 | 11 | 3 | 1 | 1, 3 |
| MY16 | 2 | 5 | 1 | 2, 4 |
| MY23 | 8 | 2 | 2 | 1, 4 |
| N2 | 4 | 1 | 5 | 2, 3 |

**Supplementary Text – Section S1: block effects**

We tested for block effects in each assay by including ‘block’ as the only predictor in a model of the defense level. In the resistance assay, block was statistically significant, so we tested further to examine block effects within each treatment and test for pairwise differences across blocks using Tukey’s Honest Significant Differences tests (functions *TukeyHSD* and *aov* in base R). In the control treatment (OP50), block 4 was significantly different from the other blocks. This block contained strain CX11314, which had very high mortality in OP50. After excluding this strain, there were no longer any block effects for the OP50 control treatment, which suggested there were not major block differences in the basic experimental protocol that would contribute to variation between strains. In the low dose Sm2170 treatment, Blocks 4 and 5 were significantly different from the other blocks, and from each other. In the high dose Sm2170 treatment, Blocks 3 and 5 were significantly different from other blocks and each other. In the low dose Db10 treatment, only Block 5 was significantly different from the others, and in the high dose Db10 treatment, there were many individual pairwise differences between blocks (2-1, 3-1, 4-2, 5-2, 4-3, 5-3), but no individual block was consistently different from the others. These block differences could be the result of the variable membership of strains in each block. In each of the main models in our analyses, we excluded data from one block at a time and ran the analysis to examine whether effects in any single block were having an outsized influence on the results, and none of the results changed qualitatively.

Next, we examined potential block effects in the avoidance assays. Block was not a statistically significant predictor of lawn-leaving. In the choice assay, block 1 had lower choice index values than the other blocks, and further tests indicated this was a function of lower avoidance of Db10 at the 21- and 27-hour timepoints in this block. Because the allocation of host strains to blocks in the choice assay was balanced, it was possible to include Block as a covariate in the models to control for it. Doing so did not change the overall effects or general conclusions of the analyses.

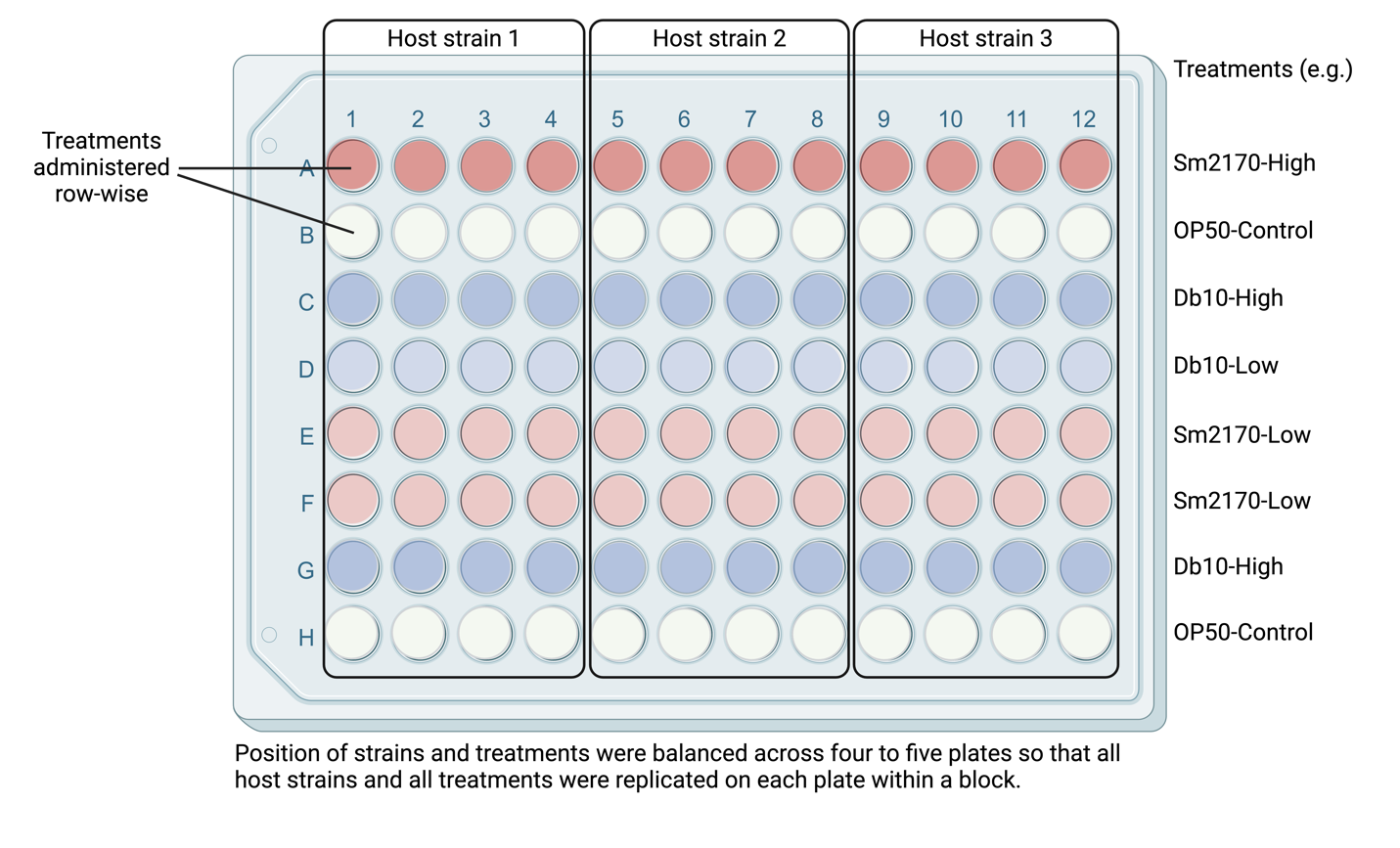

**Figure S1: Organization of host strains and treatments into 96-well plates for resistance assay**

Individual hosts from a given strain were added via the Worm Sorter to each well in four columns of a 96-well plate. Bacterial treatments (OP50 control, Sm2170 high dose, Sm2170 low dose, Db10 high dose, and Db10 low dose) were added row-wise, with each treatment represented on one or two rows per plate, for a total of six rows across all four plates. The relative positions of the treatments and the host strains were balanced across plates within a block. This resulted in a planned total of n=24 wells per host strain by treatment combination. Extra hosts were added to a fifth ‘backup’ plate so that if any wells did not contain a single live host because of Worm Sorter error, additional wells could have the corresponding doses added on the fifth plate.

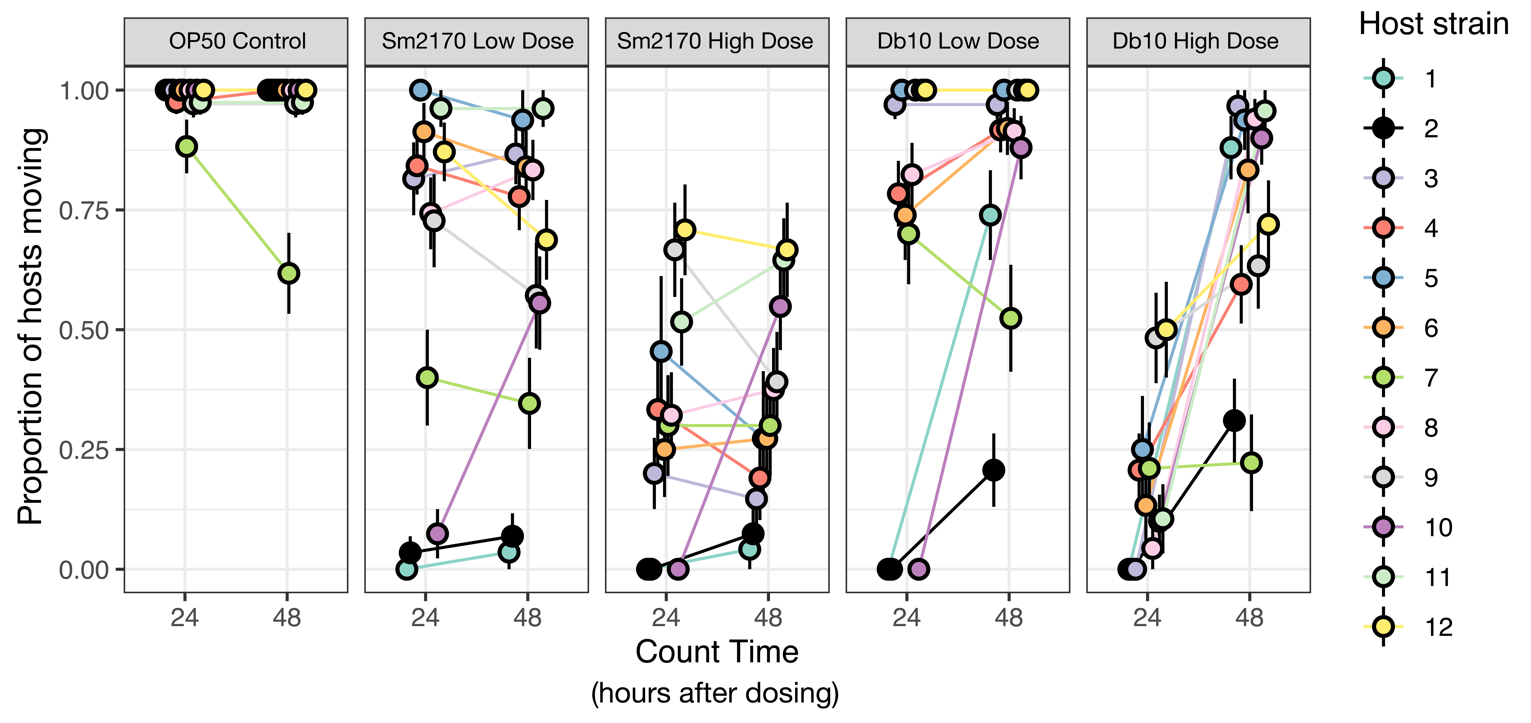

**Figure S2: Strain immobility at 24 hours**

Strains varied in their responses to dosing with two strains of *S. marcescens*. We quantified mortality as the proportion of hosts that were moving after 48 hours. We also checked hosts at 24 hours, and we expected that there would be a decrease in the proportion of hosts moving from 24 to 48 hours, as more hosts died. We expected occasional small increases in the proportion of individuals moving from 24-48 hours as a result of individual hosts being scored as missing or changes in host activity levels. However, in the Db10 high dose treatment, we observed an unexpected and widespread paralytic effect, where a large fraction of individuals moving at 48 hours had not been marked as moving at 24 hours. These differences could not be explained by differences in the number of missing worms. Strain 10 showed this paralysis at 24 hours in all *S. marcescens* treatments. We treated hosts that were moving at 48 hours as alive for our analyses, regardless of their status at 24 hours.

**Table S2: Best supported model of survival.** Analysis of deviance table for a GLMM with a binomial link where survival at 48 hours was coded as 1/0. A random effect for plate was also included. * p<0.05, ** p<0.01, *** p<0.001

| **Variable** | $\boldsymbol{\chi}^{\boldsymbol{2}}$ | **df** |
| --- | --- | --- |
| Bacterial strain | 188.02*** | 1 |
| Host strain | 282.82*** | 11 |
| Dose | 76.50*** | 1 |
| Bacterial strain x Host strain | 47.67*** | 11 |
| Bacterial strain x Dose | 0.012 | 1 |
| Host strain x Dose | 52.43*** | 11 |
| Bacterial strain x Host strain x Dose | 28.80** | 11 |

**Table S3: Best supported model of fecundity.** Analysis of deviance table for a GLMM with a binomial link where the presence of larvae at 48 hours was coded as 1/0. A random effect for plate was also included.

| **Variable** | $\boldsymbol{\chi}^{\boldsymbol{2}}$ | **df** |
| --- | --- | --- |
| Bacterial strain | 116.55*** | 1 |
| Host strain | 216.32*** | 11 |
| Dose | 266.52*** | 1 |
| Bacterial strain x Host strain | 33.37*** | 11 |
| Bacterial strain x Dose | 14.33*** | 1 |
| Host strain x Dose | 54.51*** | 11 |
| Bacterial strain x Host strain x Dose | 6.25 | 11 |

**Table S4: Best supported model of lawn-leaving.** Analysis of deviance table for a GLMM with a binomial link where the dependent variable was the counts of hosts off and on a lawn of the parasitic bacteria. A random effect for plate was also included.

| **Variable** | $\boldsymbol{\chi}^{\boldsymbol{2}}$ | **df** |
| --- | --- | --- |
| Bacterial strain | 867.64*** | 1 |
| Host strain | 111.46 *** | 11 |
| Count time | 18.84*** | 1 |
| Bacterial strain x Host strain | 95.57*** | 11 |

**Table S5: Best supported model of choice.** Analysis of deviance table for a LMM where the dependent variable was the choice index. A random effect for plate was also included.

| **Variable** | $\boldsymbol{\chi}^{\boldsymbol{2}}$ | **df** |
| --- | --- | --- |
| Bacterial strain | 5.81* | 1 |
| Host strain | 13.70 | 11 |
| Count time | 261.91*** | 2 |
| Block | 8.75* | 3 |

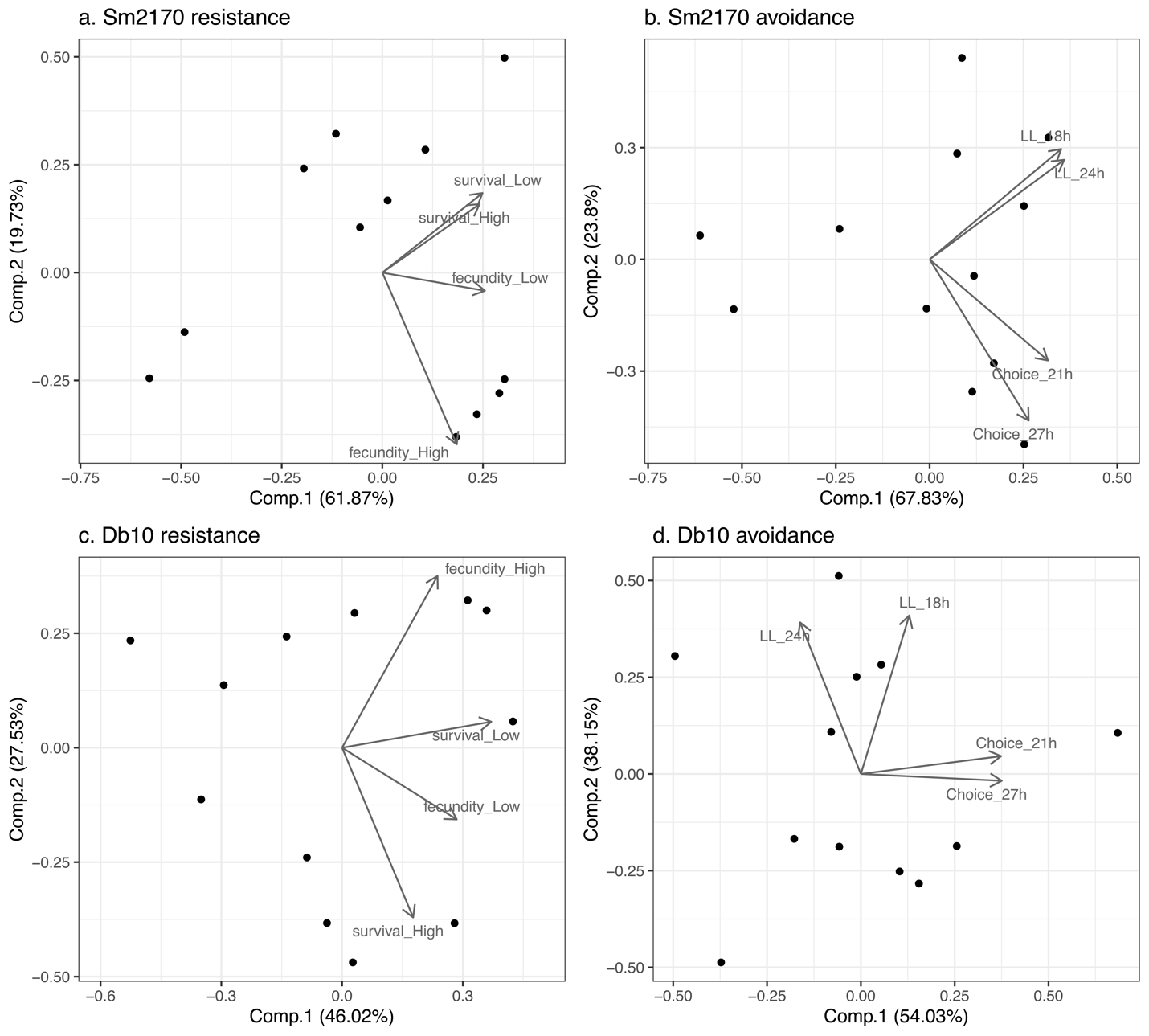

**Figure S3: Principal components analysis loadings.**

For each category of defense traits against each strain of the parasite *S. marcescens*, we ran a principal components analysis. Plotted are first two principal components for the 12 host strains (points), with the proportion of variance in the data explained by a principal component in parenthesis in the axis labels. The lines with arrowheads represent the loadings for each measure of defense level onto principal components space. The resistance traits included proportion of individuals surviving at low and high doses of the parasite (survival_Low and survival_High, respectively) and proportion of individuals reproducing at low and high doses (fecundity_Low, fecundity_High). Avoidance traits included proportion of individuals leaving the lawn of the parasite at 18 and 24 hours (LL_18h, LL_24h), as well as choice index after 21 and 27 hours (Choice_21h, Choice_27h).

**Table S6: Models testing for covariance between first principal components for resistance and avoidance.** Estimates of resistance principal component 1 from linear models predicting avoidance principal component 1. Models were fit to the two parasite strains separately. Models were not statistically significant.

| **Parasite Model** |  | **Estimate +/- SE** | **t** |
| --- | --- | --- | --- |
| Sm2170 | Intercept | 0 | 0 |
|  | Resistance PC1 | -0.31 +/-0.33 | -0.92 |
| Db10 | Intercept | 0 | 0 |
|  | Resistance PC1 | 0.31 +/-0.29 | 1.07 |
